# Powerful tests for multi-marker association analysis using ensemble learning

**DOI:** 10.1101/005405

**Authors:** Badri Padhukasahasram, Chandan K. Reddy, L. Keoki Williams

## Abstract

Multi-marker approaches are currently gaining a lot of interest in genome wide association studies and can enhance power to detect new associations under certain conditions. Gene and pathway based association tests are increasingly being viewed as useful complements to the more widely used single marker association analysis which have successfully uncovered numerous disease variants. A major drawback of single-marker based methods is that they do not consider pairwise and higher-order interactions between genetic variants. Here, we describe novel tests for multi-marker association analyses that are based on phenotype predictions obtained from machine learning algorithms. Instead of utilizing only a linear or logistic regression model, we propose the use of ensembles of diverse machine learning algorithms for constructing such association tests. As the true mathematical relationship between a phenotype and any group of genetic and clinical variables is unknown in advance and may be complex, such a strategy gives us a general and flexible framework to approximate this relationship across different sets of SNPs. We show how phenotype prediction obtained from ensemble learning algorithms can be used for constructing tests for the joint association of multiple variants. We first apply our method to simulated datasets to demonstrate its power and correctness. Then, we apply our method to previously studied asthma-related genes in two independent asthma cohorts to conduct association tests.

## INTRODUCTION

Genome wide association studies (GWAS) have generated a wealth of information about genes and genetic variants influencing various diseases and traits (Visscher 2012). The vast majority of GWAS have focused on single-marker analysis and tests for significance were corrected for multiple hypotheses testing to obtain the correct false positive rates. Because the number of markers tested in such studies is large, a single nucleotide polymorphism (SNP) needs to have strong effects or the sample size needs to be large enough to cross the stringent genome wide significance thresholds. Furthermore, many complex traits are thought to result from the interplay of multiple genetic and environmental factors, which are not captured by single SNP association tests. Given these limitations of single-marker analysis, many multi-marker approaches for association testing have been proposed and are increasingly being used to complement single SNP analysis (e.g. Pang et al. 2006; Wang et al. 2007; Li et al. 2009; Liu et al. 2010; Wu et al. 2010; Li et al. 2011; Wu et al. 2011; Huang et al. 2011; Li et al. 2012; Chung and Chen 2012).

Genes are the basic functional units of the genome and multiple polymorphisms within or near a gene can jointly affect its products. Thus, multi-marker association tests can realistically model the multiplicity that occurs biologically. While individual causal variants might show only a marginal signal of association, jointly utilizing all informative SNPs within a gene may detect their manifold effects. Testing genes also reduces the burden of multiple testing from millions of individual SNP tests to around 20,000 genes. Gene-based methods may also be less sensitive to differences in allele frequency and linkage disequilibrium between population groups (and, therefore, may produce more replicable results).

To date several gene-based association tests have been proposed (e.g. Li et al. 2009; Liu et al. 2010; Wu et al. 2010; Li et al. 2011; Wu et al. 2011; Huang et al. 2011; Li et al. 2012). Most of these approaches first assign a subset of SNPs to a particular gene based on their location in the genome; they then seek to calculate a gene-based *p* value based on the individual SNP association tests. VEGAS (Versatile gene-based association study) is a gene-based method that combines the chi-square test statistics of individuals SNPs, while accounting for their dependence (Liu et al. 2010). GATES is another gene-based association test that uses an extended Simes procedure to integrate the *p* values of individual variants while accounting for pairwise correlations between variants when calculating the effective number of independent tests (Li et al. 2011). SKAT is a logistic kernel machine based test that can account for nonlinear effects when determining the gene-level significance (Wu et al. 2010; Wu et al. 2011).

Generally, the methods used for combining *p* values in gene-based tests can be divided into 2 categories: best-SNP picking and all SNP aggregating tests. Best-SNP picking tests use only one SNP-based *p* value after accounting for multiple testing adjustment. GATES is an example of a testing method that falls within this category. All-SNP aggregating tests, such as VEGAS-SUM and SKAT, attempt to accumulate the effects of all SNPs into a test when determining the overall *p* value. HYST is a recently developed hybrid method that use both these kinds of approaches in its calculations (Li et al. 2012).

Many existing gene-based approaches either use the minimum of the *p* values for variants within a gene or integrate the *p* values/test statistics from individual variants to determine the overall gene-level *p* values. However, this may not be optimal in terms of utilizing the information available in the data and it may be better to determine the joint association of multiple predictive SNPs rather than use individual SNP measures. In addition, many existing methods do not account for nonlinear effects. Our main goal here is to develop an accurate method for multi-marker association analysis that can incorporate pairwise and higher order interactions between variables. We use phenotype prediction algorithms as a basis for constructing such association tests. Since both the underlying genetic architecture of a trait and the optimal model structure for combining the association information across multiple SNPs are not usually known before testing, we propose a machine learning approach for this purpose. The main novelty of our approach is the use of an ensemble of diverse learning models to generate phenotype predictions. In this approach, we feed the initial predictions generated from many individual learning algorithms into a second-level learning algorithm which weights their contributions suitably to generate a final prediction (Breiman 1996; Breiman 2001; Bell et al. 2007; Sill et al. 2009; Toscher et al. 2009). Thus, our approach involves blending the results of different learning algorithms by using a “meta-level” learning algorithm. We also use additional variables called “meta-features” (e.g., age, gender, body mass index, individual genotypes, ancestry) as inputs to guide this blending procedure (Sill et al. 2009). In principle, such a combination of models can allow us to better approximate (on average) the true underlying relationships between the input variables and phenotype across multiple sets of SNPs. Of note, this method allows the relationships between different groups of SNPs and the phenotype to be non-linear, complex, and variable, as is likely to occur in nature.

Here, we show how machine learning algorithms can be used to construct powerful tests for multi-marker association analysis. We then show how to construct tests of association in the presence of non-genetic covariates and how to construct a multi-marker test of interactions under this framework. We first apply our method to simulated datasets to demonstrate its power and correctness. Lastly, we apply our method to previously studied asthma-related genes in two independent asthma cohorts to conduct gene-based association tests.

## METHODS

### Approach for predicting phenotypes

Here, we present an overview of our approach to predict phenotypes from genetic and clinical variables through the use of multiple machine learning algorithms. First, we create a list of all genetic variants and clinical covariates that can potentially influence the phenotype of interest such as a disease or drug response. Next, we perform a feature selection step where we identify a subset of variables, which are useful for building a predictive model (i.e., associated with the phenotype). This can be done in many ways such as using variable importance scores from a random forest algorithm or Pearson’s correlation coefficient with the phenotype. Different machine learning algorithms (e.g., random forests (Breiman 2001), support vector machines (Cortes and Vapnik 1995, Harris et al. 1996) and logistic regression) are then trained using this subset of informative variables. Subsequently, we use the predictions from these individual models along with the selected features as inputs in a “meta-level” random forest algorithm. Lastly, we assess prediction accuracy by testing the model on an “outside the training set” and through 20-fold cross-validation.

### Ensemble learning algorithm for phenotype prediction

Ensemble learning variation 1:

1. Generate a set of all genetic variables.
2. Perform feature selection on the training data in order to identify an informative subset of variables (f_1_, f_2_…f_n_) for phenotype prediction. This can be performed using either pairwise correlation coefficients between variables and phenotype or by using random forest variable importance scores to rank the variables. Then, we can use the top 10%-30% of the variables in a prediction model.
3. Train k independent machine learning approaches on the training data using the selected features and generate model predictions P_1_, P_2_…P_k_.
4. Use the predictions from step 3, P_1_, P_2_…P_k_ and f_1_, f_2_…f_n_ as inputs and train a “meta-level” learning algorithm using random forests. Note that this is a key step in the algorithm and generates a final prediction by blending many individual predictions in a possibly nonlinear manner. The main goal is to learn the best model to combine individual models from the training data so that we can predict the phenotype as well as possible. The non-linear combination of models along with the meta-features gives us a more general predictive framework, which can accommodate different model structures and also allows the overall model to vary across the multi-dimensional parameter space.
5. Generate predictions in test data P_blend1_ using the models trained in steps 3 and then 4. Repeat for all cross-validation folds to obtain unbiased phenotype predictions for all samples.

### Generalization: An ensemble of ensembles

Generalizations of the algorithm described previously are also possible that can potentially further boost the prediction accuracy. In particular, the creation of an ensemble of models (steps 3 and 4 in previous algorithm) can be done in a variety of different ways. For example:

Ensemble learning variation 2: Combining of predictions from individual learning models can be done sequentially using predictions from all previous steps as inputs in the next step (i.e. instead of steps 3 and 4). Therefore, as an alternative approach, we can do the following:

i. Train learning algorithm 1 on the training data using the selected features f_1_, f_2_…f_n_ as inputs and generate model predictions P_1_.
ii. Train learning algorithm 2 on the training data using P_1_ and the selected features f_1_, f_2_…f_n_ as inputs and generate model predictions P_2_.
iii. Training learning algorithm 3 on the training data using P_1_, P_2_ and the selected features f_1_, f_2_…f_n_ as inputs and generate model predictions P_3_.

………………………………………………………………………………………………..……

…………………………………………………………………………………………………..…

k) Training learning algorithm k on the training data using P_1_, P_2_,…P_k-1_ and the selected features f_1_, f_2_…f_n_ as inputs and generate model predictions P_k_.

Note that each algorithm after the first is a meta-level learning algorithm. Then, we generate predictions in test data P_blend2_ using the models as in training and repeat for all cross-validation folds to obtain unbiased phenotype predictions for all samples.

Ensemble learning variation 3: Instead of applying an ensemble learning model (variation 1) to all the samples, we can divide the high-dimensional parameter space of variables into different subsets. Then, we can train different ensemble learning models using only samples that fall in these different subsets and finally merge these models to obtain the overall prediction model. Subsequently, we can generate final predictions, P_blend3_, in test data as we did for training data for all cross-validation folds within all subsets to obtain unbiased phenotype predictions for the entire sample.

Lastly, we can train a final random forest learning algorithm that uses f_1_, f_2_…f_n_ and P_blend1_, P_blend2_ and P_blend3_ as inputs and performs 20-fold cross-validation to generate the final prediction P_final_.

### Multi-marker tests of association

Once we have estimated a model using any of the algorithms described in the previous section and predicted phenotypes for all individuals using cross-validation, we can construct tests of association in the following manner. For continuous traits, we can calculate the Pearson’s correlation coefficient between predicted (P_final_) and observed (P_actual_) values and obtaining the corresponding *p* values. For case-control studies, we perform a logistic regression using all the genetic variables (i.e., SNPs) and P_final_ as explanatory variables. A chi square based likelihood ratio test can then be used to generate *p* values.

### Testing multi-marker associations in the presence of covariates

Association testing in the presence of covariates (e.g., age, gender, BMI and smoking status) can be done in the following manner. First, consider both non-genetic covariates and genetic variables together for phenotype prediction according to any of the ensemble learning algorithms described earlier. Let P_final-all_ be the predicted phenotype values. Then, remove the SNP variables and rerun the phenotype prediction algorithm. Let P_final-covariates_ be the predicted phenotype values. For continuous traits, we first calculate the Pearson’s correlation coefficient for these predicted variables with the true phenotypes (P_actual_). The strength of association for the genetic variables can then be calculated using the Steiger’s *Z* test (Steiger 1980) for the difference between the 2 calculated correlation coefficients. Let *r_12_* and *r_13_* denote the Pearson’s correlations between the true phenotype (P_actual_) and P_final-covariates_ and P_final-all_ respectively. Let *r_23_* denote the Pearson’s correlation between P_final-covariates_ and P_final-all_. The Steiger’s test computes *p* values based on the following test statistic that is assumed to be standard normally distributed:

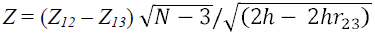

Here, *Z_12_* and *Z_13_* are Fisher’s transformations of *r_12_* and *r_13_*, and

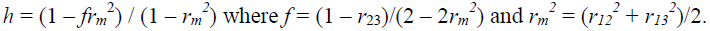

For case-control studies we can use both non-genetic covariates, genetic variables, P_final-all_, and P_final-covariates_ as explanatory variables in a logistic regression model. We then use a chi square based likelihood ratio test to compare the former model with a model without any genetic variables (i.e. non-genetic covariates and P_final-covariates_ only) to calculate a *p* value for the genetic contribution.

### Multi-marker tests for interactions

We can test for interactions between a set of markers in the following manner. First, consider all of the SNPs together in a linear or logistic regression model (for continuous or case-control phenotype) and generate phenotype predictions using cross-validation for all individuals. Let P_linear_ be the predicted phenotype values. Then, generate phenotype predictions for all individuals using any of the ensemble learning algorithms described previously. Let P_ensemble_ denote the predicted phenotype values. For continuous traits, we will use all markers as well as P_ensemble_ and P_linear_ as explanatory variables in a multiple regression model (Model 1) and perform a F test with a model (Model 0) without interactions (i.e. one with all markers and P_linear_ only) to calculate the *p* value. We compare the sum of the squared errors (SSE) of prediction to construct an F statistic with (1, N – V_Model1_ – 1) degrees of freedom. Here: F = [SSE_Model0_ – SSE_Model1_][N – V_Model1_ - 1]/SSE_Model1_. N denotes the number of samples and V_Model1_ denotes the total number of explanatory variables in model 1. For case-control studies, we will use all markers as well as P_ensemble_ and P_linear_ as explanatory variables in a logistic regression model and use a chi square based likelihood ratio test with a model without interactions (i.e. one with all markers and P_linear_ only) to calculate the *p* value.

### Power and Type-1 error rates of gene-based association tests for data simulated under multiplicative and additive models

We tested the performance of the proposed gene-based test by simulating genotype data for 30 biallelic SNPs assuming Hardy Weinberg equilibrium. We assumed the following 3 scenarios of linkage disequilibrium (LD) for the 30 SNPs: i) SNPs are within blocks with high LD (r = 0.9 or 0.8 within blocks); ii) SNPs are within blocks in moderate LD (r = 0.5 or 0.4); and iii) SNPs are completely independent of one another and in linkage equilibrium. The choice of simulation settings were similar to what has been used previously (Li et al. 2011). For each LD scenario, we considered 3 different gene sizes with the first 3, first 10 and all 30 SNPs with 1, 2 and 6 causative SNPs respectively. For each gene size, we tested the following models: i) a null model with no disease loci ii) an additive model where one SNP in each LD block had a minor allele that increased the risk additively by 0.14; and iii) a multiplicative model where one SNP in each LD block had a minor allele that increased the risk by a factor of 1.14. Disease prevalence was assumed to be 0.1. For each scenario, we used a sample of 1,500 cases and 1,500 controls drawn from a simulated population of 100,000 individuals. More details about LD patterns can be found elsewhere (Li et al. 2011). Type-1 error rates and statistical power were obtained from 1,000 and 500 simulated case-control datasets, respectively and were based on the fraction of datasets for which the gene-based association test generated significant *p* values (i.e. *p* < 0.05).

### Power and Type-1 error rates of a gene-based test for models with interactions

The simulations in the previous section assumed that the effect of various disease susceptibility SNPs were independent of one another and that they increased the risk additively or multiplicatively. To explore the effect of pairwise and higher order interactions between genetic variants, we also compared the performance of methods for data simulated under models with interactions. We simulated a quantitative trait for many different models with one or more interactions among variants in addition to main effects. In addition, we also considered scenarios where there is pure epistasis (i.e. where the effect of a group of SNPs is simply due to their interactions and there are no main effects). We simulated samples of 3,000 individuals and genes with 5 or 10 SNPs assuming linkage equilibrium. The phenotype was drawn from a complex distribution involving the sum of a standard normal random variable and some multivariable function involving many SNP variables (Table 4). SNP variables are coded as 0, 1 or 2. Power and Type-1 error rates were estimated based on 100 and 500 simulated datasets, respectively. We calculated the fraction of simulated datasets for which the gene-based method generated a significant *p* value (*p* < 0.05). We compared our result with a gene-based test using linear regression and a gene-based test using GATES (Li et al. 2011). For the gene-based test with linear regression, *p* values were obtained using an F test statistic.

### Power and Type-1 Error rates for a multi-marker test for interactions

For all the models simulated in the previous section, we also constructed a multi-marker test for interactions as described previously and estimated the power of such a test. We simulated samples of 3,000 individuals and genes with 5 or 10 SNPs assuming linkage equilibrium. The phenotype was drawn from a complex distribution involving a sum of a standard normal variable and interaction terms involving many SNPs as shown in Table 4. Power and type-1 error rates were estimated based on 1,000 simulated datasets. For each model with interactions, we calculated the fraction of simulated datasets for which the multi-marker test of interactions generated a significant *p* value (*p* < 0.05); *p* values were based on an F test statistic with two parameters as described previously.

### Datasets

We applied the methods developed in this paper to data from 2 independent studies. The studies included the Study for Asthma Phenotypes and Pharmacogenomic Interactions by Race-ethnicity (SAPPHIRE) and the Genes-environments and Admixture in Latino Americans (GALA II). Recruitment for both studies is ongoing.

SAPPHIRE is population-based study which seeks to understand the genetic underpinnings of both asthma and asthma medication response. Study individuals included in this analysis were recruited from a single large health system serving the southeast Michigan and the Detroit metropolitan area. Enlisted patients with asthma met the following criteria: age 12-56 years, a physician diagnosis of asthma, and no recorded diagnosis of chronic obstructive pulmonary disease or congestive heart failure. Control individuals without asthma were recruited from a similar geographic region and were 12-56 years of age, but they did not have a prior recorded diagnosis of asthma, chronic obstructive pulmonary disease, or congestive heart failure. Genome wide genotyping was performed using the Axiom Genome-Wide AFR array (Affymetrix, Santa Clara, CA). After data quality control, genotype information was available on 586,952 SNPs for 1,073 individuals with asthma and 328 healthy controls (Padhukasahasram et al. 2014). All of the individuals from the SAPPHIRE cohort included in this analysis were African American by self-report.

The GALAII study is a case control study to identify gene-environment interactions contributing to asthma. Children of Latino descent age 8-21 years were recruited from New York City, Chicago, San Francisco, Houston, and Puerto Rico. Children with asthma had a physician diagnosis of asthma and either a 12% increase in forced expiratory volume at one second following the administration of albuterol or a positive methacholine challenge test. Genome wide genotype data was available on 3,772 individuals (1,891 with asthma and 1,881 without). Genomic DNA was genotyped on the Axiom Genome-Wide LAT array. After data cleaning, information was available for 747,075 SNPs genome wide.

## RESULTS

### Multiplicative and Additive models-Comparisons

Tables 1–3 shows comparisons for the performance of various methods for disease case-control datasets simulated under additive and multiplicative models. We can see that the performance of the newly proposed method based on an ensemble of machine learning algorithms is comparable to other approaches and the Type-1 error rates produced by all methods are close to what is expected. For more details about the different methods tested in these tables, please refer to Li et al. 2011. Note that when there are no disease-related SNPs in the data, we expect to see *p* values < 0.05, in around 5% of the simulated datasets due to chance alone. For the ensemble learning and logistic regression methods, we can also see that power is not strongly sensitive to the strength of linkage disequilibrium. Thus, for both additive and multiplicative models, power estimates do not appear to change much across Tables 1-3 for these methods.

**Table 1.** Comparison of empirical power and Type 1 error rates of gene-based association tests for simulated datasets assuming linkage equilibrium.

|  | #SNP<br>(#DSL) | Logistic<br>Regression | Fisher | Vegas-Sum | Original<br>-Simes | Vegas-Max | GATES | Machine-Learning<br>Ensemble |
| --- | --- | --- | --- | --- | --- | --- | --- | --- |
| Linkage Equilibrium |  |  |  |  |  |  |  |  |
| Type 1 Error | 3(0) | 4.66 | 4.67 | 4.70 | 4.61 | 4.62 | 4.61 | 4.90 |
| Type 1 Error | 10(0) | 5.10 | 5.00 | 5.04 | 5.06 | 5.07 | 5.06 | 4.80 |
| Type 1 Error | 30(0) | 5.26 | 4.96 | 4.97 | 4.97 | 5.04 | 4.97 | 5.60 |
| Power<br>Additive | 3(1) | 43.71 | 41.79 | 42.67 | 45.28 | 45.22 | 45.28 | 56.00 |
| Power<br>Additive | 10(2) | 56.88 | 53.32 | 54.56 | 54.76 | 54.00 | 54.76 | 57.60 |
| Power<br>Additive | 30(6) | 65.32 | 61.5 | 63.28 | 47.18 | 45.62 | 47.18 | 69.00 |
| Power<br>Multiplicative | 3(1) | 46.61 | 44.72 | 45.54 | 48.39 | 48.3 | 48.39 | 53.00 |
| Power<br>Multiplicative | 10(2) | 69.00 | 65.25 | 66.88 | 67.00 | 66.26 | 67.00 | 69.00 |
| Power<br>Multiplicative | 30(6) | 93.45 | 91.44 | 92.28 | 82.21 | 80.18 | 82.21 | 94.60 |

DSL denotes the number of disease susceptibility markers.

**Table 2.** Comparison of empirical power and Type 1 error rates of gene-based association tests in simulated datasets for moderate linkage disequilibrium.

|  | #SNP<br>(#DSL) | Logistic<br>Regression | Fisher | Vegas-Sum | Original<br>-Simes | Vegas-Max | GATES | Machine-Learning<br>Ensemble |
| --- | --- | --- | --- | --- | --- | --- | --- | --- |
| Linkage Disequilibrium |  |  |  |  |  |  |  |  |
| Type 1 Error | 3(0) | 4.86 | 7.17 | 4.91 | 4.54 | 4.81 | 4.98 | 5.20 |
| Type 1 Error | 10(0) | 4.88 | 9.8 | 4.83 | 4.55 | 4.92 | 5.00 | 5.60 |
| Type 1 Error | 30(0) | 5.63 | 11.09 | 5.03 | 4.97 | 5.29 | 5.56 | 5.70 |
| Power<br>Additive | 3(1) | 44.59 | 55.8 | 49.36 | 49.71 | 50.51 | 51.23 | 55.20 |
| Power<br>Additive | 10(2) | 56.25 | 72.38 | 61.36 | 58.39 | 59.12 | 60.72 | 63.80 |
| Power<br>Additive | 30(6) | 65.47 | 83.04 | 71.96 | 53.29 | 52.24 | 55.65 | 68.00 |
| Power<br>Multiplicative | 3(1) | 46.52 | 57.5 | 50.98 | 51.19 | 52.00 | 52.65 | 53.40 |
| Power<br>Multiplicative | 10(2) | 68.42 | 81.73 | 72.48 | 70.66 | 70.9 | 72.4 | 70.20 |
| Power<br>Multiplicative | 30(6) | 93.68 | 98.04 | 95.59 | 86.07 | 84.34 | 87.52 | 94.70 |

DSL denotes the number of disease susceptibility markers.

**Table 3.** Comparison of empirical power and Type 1 error rates of gene-based association tests on simulated datasets for strong linkage disequilibrium.

|  | #SNP<br>(#DSL) | Logistic<br>Regression | Fisher | Vegas-Sum | Original<br>-Simes | Vegas-Max | GATES | Machine-Learning<br>Ensemble |
| --- | --- | --- | --- | --- | --- | --- | --- | --- |
| Linkage Disequilibrium |  |  |  |  |  |  |  |  |
| Type 1 Error | 3(0) | 4.96 | 11.49 | 5.23 | 3.88 | 5.22 | 5.35 | 6.00 |
| Type 1 Error | 10(0) | 5.33 | 15.68 | 4.84 | 3.37 | 4.88 | 5.34 | 5.70 |
| Type 1 Error | 30(0) | 5.57 | 17.9 | 4.89 | 3.38 | 4.89 | 5.64 | 5.90 |
| Power<br>Additive | 3(1) | 45.03 | 72.29 | 58.81 | 53.88 | 58.2 | 60.43 | 61.00 |
| Power<br>Additive | 10(2) | 57.20 | 89.82 | 75.74 | 66.39 | 71.71 | 74.3 | 59.00 |
| Power<br>Additive | 30(6) | 65.56 | 96.04 | 86.3 | 62.84 | 66.80 | 72.75 | 65.80 |
| Power<br>Multiplicative | 3(1) | 47.13 | 74.28 | 60.88 | 56.28 | 60.74 | 62.77 | 65.00 |
| Power<br>Multiplicative | 10(2) | 68.45 | 94.41 | 84.89 | 77.14 | 80.59 | 83.00 | 74.40 |
| Power<br>Multiplicative | 30(6) | 93.4 | 99.92 | 99.2 | 91.42 | 92.24 | 95.38 | 94.00 |

DSL denotes the number of disease susceptibility markers.

### Models with epistatic effects

In Table 4, we show the power of the ensemble learning based multi-marker association test using a simulated quantitative trait for models with interactions. We compare the ensemble learning approach with a gene-based test constructed using multiple linear regression, as well as with the extended Simes procedure as implemented by GATES. In all situations, our simulations indicate that the machine learning approach, which can model interactive effects, is uniformly more powerful for detecting gene-based associations when compared with the other two approaches. Table 4 also shows that the estimated gain in power can be substantial. Among the other two methods, multiple linear regression performed second best while the GATES method which only integrates the *p* values from single marker tests had the lowest power. In Table 5, we show the power and Type-1 error rates of a multi-marker test for interactions using the same models as in Table 4. These results clearly demonstrate the ability of our approach to detect the presence of interactions by considering the difference between ensemble learning and linear model based predictions.

**Table 4.** Comparison of empirical Power and Type-1 error rates of gene-based association tests for a quantitative trait simulated under models with interactions.

| Value | Phenotype distribution | #SNP<br>(#TAS) | Linear<br>Regression | GATES | Machine<br>learning<br>Ensemble |
| --- | --- | --- | --- | --- | --- |
| Type 1 error | $P \sim N(0,1)$ | 5(0) | 5.66 | 4.00 | 5.00 |
| Type 1 error | $P \sim N(0,1)$ | 10(0) | 5.30 | 6.00 | 5.66 |
| Power | $P \sim N(0,1) + 0.20 * snp1 * snp2 * snp9 * snp10$ | 10(4) | 40.0 | 36.0 | 44.0 |
| Power | $P \sim N(0,1) + 0.002 * snp1 + 0.002 * snp2 + 0.12 * snp1 * snp2 + 0.18 * snp3 * snp4$ | 5(4) | 84.0 | 66.0 | 87.0 |
| Power | $P \sim N(0,1) + 0.25 * snp1 * snp2 * snp3$ | 5(3) | 38.0 | 34.0 | 56.0 |
| Power | $P \sim N(0,1) + 0.3 * snp1 * snp2 * snp3$ | 5(3) | 49.0 | 36.0 | 64.0 |
| Power | $P \sim N(0,1) + 0.35 * snp2 * snp3 * snp4$ | 5(3) | 64.0 | 55.0 | 87.0 |
| Power | $P \sim N(0,1) + 0.65 * snp1 * snp2 * snp3 * snp8 * snp9 * snp10$ | 10(6) | 58.0 | 47.0 | 85.0 |
| Power | $P \sim N(0,1) + 0.002 * snp1 + 0.002 * snp2 + [0.2 * (1 + snp1) / (1 + snp2)] + 0.3 * snp4 * snp5$ | 5(4) | 66.0 | 50.0 | 80.0 |
| Power | $P \sim N(0,1) + 0.002 * snp1 + 0.002 * snp2 + 0.3 * snp1 * snp2 + 0.2 * snp3 * snp4$ | 5(4) | 61.0 | 49.0 | 96.0 |

TAS denotes the number of trait associated SNPs.

**Table 5.** Empirical Power and Type-1 error rate of a gene-based test of interactions for a simulated quantitative trait.

| Value | Phenotype distribution | #SNP<br>(#TAS) | Machine<br>learning<br>Ensemble |
| --- | --- | --- | --- |
| Type 1 error | $P \sim N(0,1)$ | 5(0) | 6.10 |
| Type 1 error | $P \sim N(0,1)$ | 10(0) | 5.10 |
| Power | $P \sim N(0,1) + 0.20 * \text{snp1} * \text{snp2} * \text{snp9} * \text{snp10}$ | 10(4) | 55.5 |
| Power | $P \sim N(0,1) + 0.002 * \text{snp1} + 0.002 * \text{snp2} + 0.12 * \text{snp1} * \text{snp2} + 0.18 * \text{snp3} * \text{snp4}$ | 5(4) | 52.5 |
| Power | $P \sim N(0,1) + 0.25 * \text{snp1} * \text{snp2} * \text{snp3}$ | 5(3) | 64.6 |
| Power | $P \sim N(0,1) + 0.3 * \text{snp1} * \text{snp2} * \text{snp3}$ | 5(3) | 78.2 |
| Power | $P \sim N(0,1) + 0.35 * \text{snp2} * \text{snp3} * \text{snp4}$ | 5(3) | 87.5 |
| Power | $P \sim N(0,1) + 0.65 * \text{snp1} * \text{snp2} * \text{snp3} * \text{snp8} * \text{snp9} * \text{snp10}$ | 10(6) | 87.4 |
| Power | $P \sim N(0,1) + 0.002 * \text{snp1} + 0.002 * \text{snp2} + [0.2 * (1 + \text{snp1}) / (1 + \text{snp2})] + 0.3 * \text{snp4} * \text{snp5}$ | 5(4) | 53.5 |
| Power | $P \sim N(0,1) + 0.002 * \text{snp1} + 0.002 * \text{snp2} + 0.3 * \text{snp1} * \text{snp2} + 0.2 * \text{snp3} * \text{snp4}$ | 5(4) | 83.6 |

TAS denotes the number of trait associated SNPs.

### Application to real datasets

We applied the proposed gene-based association test to an empirical dataset consisting of 1,401 African American individuals (1,073 individuals with asthma and 328 individuals without asthma) from the SAPPHIRE cohort and 3,772 Latino children (1,891 individuals with asthma and 1,881 individuals without asthma) from the GALAII study. Tables 6 and 7 show the sample characteristics of these populations. We tested 9 previously studied asthma-related genes (Li et al. 2010; Moffatt et al. 2010; Torgerson et al. 2011) to see if these are also associated with asthma status in our datasets. Although 100s of genes have been implicated in asthma, only a few have been reliably replicated in multiple groups. Therefore, to demonstrate the performance of our method, we restricted our analysis to a small subset of asthma genes identified (and replicated in some cases) in well-powered, high-quality studies. This also reduces the burden of multiple testing. When constructing gene-based association tests, we adjusted for age, gender and the first 10 principal components in both study groups. Principal component analysis was performed using the prcomp function in R using a random set of 10,000 markers. Tables 8 and 9 show the results of our ensemble learning gene-based association test in the SAPPHIRE and GALAII study populations, respectively. The results are compared with those obtained using the GATES method and logistic regression. At a Bonferroni adjusted significance threshold of 0.0027 (= 0.05/18 [i.e., 9 genes assessed twice]), we found that the ensemble learning gene-based test identified more statistically significant results when compared with the other gene-based methods. Specifically, IL33 was significantly associated with asthma in Latino children using the ensemble learning gene-based test, but this gene was of borderline significance using the other 2 approaches.

**Table 6.** Sample characteristics of the SAPPHIRE cohort.

| **Variable** | **Healthy African American**<br>**individuals**<br>**(n=328)** | **African American individuals**<br>**with asthma**<br>**(n=1,073)** |
| --- | --- | --- |
| Age (years) – mean ± SD | 41.23 ± 13.28 | 31.65 ± 14.57 |
| Female Sex – no. (%) | 212 (64.63) | 671 (62.53) |
| Body mass index (kg/m<sup>2</sup>) –<br>mean ± SD | 32.19 ± 7.58 | 31.49 ± 9.07 |
| Smoking status – no. (%) |  |  |
| Never | 239 (72.8) | 893 (83.2) |
| Past | 33 (10.1) | 96 (8.9) |
| Current | 56 (17.1) | 84 (7.8) |
| Asthma age of onset (years)<br>– mean ± SD | -- | 12.65 ± 13.55 |
| FEV<sub>1</sub> (liters) – mean ± SD | 2.74 ± 0.71 | 2.58 ± 0.75 |
| Percent of predicted FEV<sub>1</sub> –<br>mean ± SD | 97.6 ± 15.3 | 87.9 ± 18.4 |
| SABA response (% change)<br>– mean ± SD* | 2.51 ± 7.95 | 10.53 ± 12.93 |

**Table 7.** Sample characteristics of the GALAII cohort.

| **Variable** | **Healthy Latino children**<br>**(n=1,881)** | **Latino children with asthma**<br>**(n=1,891)** |
| --- | --- | --- |
| Age (years) – mean ± SD | 13.65 ± 3.50 | 12.53 ± 3.25 |
| Female Sex – no. (%) | 1059 (56.29%) | 845 (44.68%) |
| Body Mass index (kg/m<sup>2</sup>)<br>mean ± SD | 24.40 ± 6.87 | 23.09 ± 6.51 |
| Smoking status- no(%) |  |  |
| Smoker | 92 (4.89%) | 57(3.01%) |
| Non-smoker | 1789 (95.11%) | 1834(96.99%) |
| Ethnicity – no(%) |  |  |
| Mexican | 661 (35.14%) | 596 (31.52%) |
| Peurto Rican | 894 (47.53%) | 892 (47.17%) |
| Spanish | 125 ( 6.64%) | 244 (12.90%) |
| Other | 201 (10.69%) | 159 ( 8.41%) |

**Table 8.** Gene-based *p* values for previously reported asthma-related genes in 1,401 African American individuals from the SAPPHIRE cohort.

| Chromosome | Gene | Length in<br>base pairs | Number of<br>SNPs<br>tested | Gene-based<br>$p$ value<br>from<br>Ensemble<br>Learning | Gene-based<br>$p$ value<br>from<br>Logistic<br>Regression | Gene-based<br>$p$ value<br>from<br>GATES |
| --- | --- | --- | --- | --- | --- | --- |
| 1 | PYHIN1 | 45513 | 13 | 0.198 | 0.230 | 0.130 |
| 2 | IL1RL1 | 7466 | 6 | 0.982 | 0.982 | 0.832 |
| 5 | TSLP | 6333 | 5 | 0.063 | 0.064 | 0.533 |
| 9 | IL33 | 42198 | 12 | 0.408 | 0.180 | 0.130 |
| 17 | GSDMB | 14056 | 15 | 0.401 | 0.401 | 0.870 |
| 5 | IL13 | 2937 | 3 | 0.156 | 0.164 | 0.387 |
| 15 | SMAD3 | 57175 | 23 | 0.323 | 0.323 | 0.359 |
| 5 | SLC22A5 | 25906 | 15 | 0.0095 | 0.162 | 0.0076 |
| 5 | RAD50 | 87698 | 34 | 0.010 | 0.010 | 0.367 |

**Table 9.** Gene-based *p* values for previously reported asthma-related genes in 3,772 Latino individuals from the GALA study.

| Chromosome | Gene | Length<br>in<br>base pairs | Number<br>of SNPs<br>tested | Gene-based<br>$p$ value<br>from<br>Ensemble<br>Learning | Gene-based<br>$p$ value<br>from<br>Logistic<br>Regression | Gene-based<br>$p$ value<br>from<br>GATES |
| --- | --- | --- | --- | --- | --- | --- |
| 1 | PYHIN1 | 45513 | 15 | 0.320 | 0.320 | 0.530 |
| 2 | IL1RL1 | 7466 | 16 | 0.038 | 0.038 | 0.046 |
| 5 | TSLP | 6333 | 7 | 0.270 | 0.270 | 0.250 |
| 9 | IL33 | 42198 | 14 | 0.0014 | 0.095 | 0.069 |
| 17 | GSDMB | 14056 | 13 | 2.33E-09 | 4.20E-08 | 6.24E-11 |
| 5 | IL13 | 2937 | 10 | 0.280 | 0.280 | 0.100 |
| 15 | SMAD3 | 57175 | 28 | 0.464 | 0.464 | 0.063 |
| 5 | SLC22A5 | 25906 | 12 | 0.838 | 0.838 | 0.956 |
| 5 | RAD50 | 87698 | 33 | 0.217 | 0.217 | 0.050 |

## DISCUSSION

We have introduced a new method for assessing gene-based associations using genome wide genotype data. This method uses diverse machine learning algorithms to construct predictive models for the phenotype using the SNP variation with a gene and then using these predictions to construct tests of association. Machine learning algorithms represent powerful tools for inferring the relationship between multiple explanatory variables and a phenotype while accounting for complicated interactions between the former. Because the “true” multivariable relationship between a set of variables and a trait like disease or drug response is not known in advance, we can better approximate this relationship by first learning from the data. The use of ensemble learning-based predictions leads to novel multi-marker tests of association. In addition to gene-based tests of association, we expect that these methods could also be applied for pathway-based analysis or to any other set of polymorphic variants defining a region of interest or a functional class.

There are three key advantages of using our gene-based approach compared to existing approaches. First, our method does not make *a priori* assumptions about the genetic model for a SNP (i.e. additive, recessive or dominant). When constructing our tests, we can include 3 variables for each SNP where the variants are encoded according to these 3 models (i.e. additive, recessive, dominant). Thus, we can include heterogeneous effects within and across SNPs. A second advantage is the ability to include any number of covariates (genetic or non-genetic) and model higher level interactions between them. This feature makes machine learning particularly suited for assessing gene-environment or gene-gene interactions. Third, creating an ensemble of diverse multivariate models with meta-features makes our method less restrictive and capable of approximating the phenotype more accurately. Collectively, these novel aspects can boost statistical power and result in novel genetic discoveries.

Extensions of these methods towards the case of multiple correlated phenotypes should also be straightforward. If instead of a single phenotype, we are interested in many phenotypes that are correlated with one another in some manner, we can construct a joint association test for all of them in the following manner. First, we will apply the ensemble learning based gene-based association test to each phenotype individually and obtain their corresponding *p* values. Subsequently, we can obtain an overall *p* value from these individual *p* values using the TATES multi-trait association method (van der et al. 2013), which is analogous to the extended Simes procedure of GATES developed for testing multi-marker associations.

We applied our method to both simulated and empirical datasets to demonstrate its power and utility. For models without interactions between variables, the ensemble learning approach worked similarly when compared with other previous gene-based association tests. In contrast, for models dominated by interactions, our simulation studies suggested that the ensemble learning test can be considerably more powerful than other methods. Thus, for situations where epistatic or gene-environment effects are likely to be important, our association test is more likely to detect associations as compared to the alternative methods described.

There are a number of potential limitations to our approach that require mentioning. First, computational time can be a limitation when applying an ensemble learning algorithm based associations tests to thousands of genes. One potential solution would be to start by using a computationally efficient gene-based method, such as the GATES procedure, to first identify a smaller subset of likely candidate genes. Then, a machine learning based multi-marker association approach could be applied to this restricted set to further refine the group of candidate genes. However, at this point it is uncertain whether such an approach would result in improved statistical power. Next, we cannot state with certainty that the genes assessed here are involved in asthma pathogenesis, since many of these genes were identified in association studies and their function (as it relates to asthma) has not yet been elucidated. Therefore, while we assume that these genes represent true-positives, this portion of our analysis may not represent an actual demonstration of statistical power unless more detailed functional studies are conducted for the relevant genes to directly demonstrate their role in asthma. Lastly, it should also be mentioned that while our multi-marker tests can detect associations or the presence of interactive effects, they do not attempt to pinpoint the specific variants contributing to such effects. Elucidating such details will entail more in-depth analysis of the models learned and construction of additional tests.

In summary, ensemble learning algorithms provide a general and flexible framework for conducting association analysis. We have shown that phenotype predictions made by such algorithms can be used for many common tasks encountered in association analysis, such as multi-marker association tests, adjusting for genetic and non-genetic covariates, and tests of interaction. Because machine learning is a highly developed area of study, prediction of response from many input variables is a well-studied problem and numerous well-established algorithms are already available which can be readily incorporated as components in an ensemble learning framework to maximize prediction accuracy and construct powerful tests of association.

